# Cell type-specific structural plasticity of the ciliary transition zone in *C. elegans*

**DOI:** 10.1101/382689

**Authors:** Jyothi S. Akella, Malan Silva, Natalia S. Morsci, Ken C. Nguyen, William J. Rice, David H. Hall, Maureen M. Barr

**Affiliations:** Department of Genetics and Human Genetics Institute of New Jersey, Rutgers University, Piscataway, NJ 08854, USA; Department of Biology, University of Utah, Salt Lake City, UT 84112, USA; Center for *C. elegans* Anatomy, Albert Einstein College of Medicine, Bronx, NY 10461, USA; Simons Electron Microscopy Center, New York Structural Biology Center, NY 10027, USA; Waksman Institute for Microbiology, Rutgers University, Piscataway, NJ 08854, USA

**Keywords:** cilium, ciliogenesis, neuron, transition zone, microtubule, remodeling

## Abstract

**Background information:** The current consensus on cilia development posits that the ciliary transition zone (TZ) is formed via extension of nine centrosomal microtubules. In this model, TZ structure remains unchanged in microtubule number throughout the cilium life cycle. This model does not however explain structural variations of TZ structure seen in nature, and could also lend itself to the misinterpretation that deviations from nine-doublet microtubule ultrastructure represent an abnormal phenotype. Thus, a better understanding of events that occur at the TZ *in vivo* during metazoan development is required.

**Results:** To address this issue, we characterized ultrastructure of two types of sensory cilia in developing *Caenorhabditis elegans*. We discovered that, in cephalic male (CEM) and inner labial quadrant (IL2Q) sensory neurons, ciliary TZs are structurally plastic and remodel from one structure to another during animal larval development. The number of microtubules doublets forming the TZ can be increased or decreased over time, depending on cilia type. Both cases result in structural TZ intermediates different from TZ in adult cilia. In CEM cilia, axonemal extension and maturation occurs concurrently with TZ structural maturation.

**Conclusions and Significance:** Our work extends the current model to include the structural plasticity of metazoan transition zone, which can be structurally delayed, maintained or remodeled in cell type-specific manner.

## Introduction

The cilium is a conserved eukaryotic organelle present in most non-dividing cells of the human body. Cilia structure has profound importance for their functionality and functionality of the nascent cells. All cilia and flagella share common structure consisting of a microtubule-based core, called the axoneme, surrounded by a ciliary membrane. The most widespread axonemal structure consists of nine doublet microtubules (dMTs) arranged in a circle in radial symmetry around a pair of singlet microtubules (sMTs) (annotated as 9+2 motile or 9+0 sensory/primary cilia, depending on presence of inner microtubules). The proximal-most section of ciliary axoneme is the transition zone (TZ), identified by the presence of multiple rows of Y-shaped links connecting axonemal dMTs to the ciliary membrane. The ciliary TZ is the physical location of a diffusion barrier that separates the ciliary compartment from the rest of the cell and modulates ciliary composition and function (Chih et al., 2011; Garcia-Gonzalo et al., 2011; Jensen et al., 2015; Reiter et al., 2012; Takao and Verhey, 2016).

The term ciliogenesis refers to multi-step process of cilium/flagellum formation starting with basal body maturation up to formation of its functional axoneme (Ishikawa and Marshall, 2011; Sorokin, 1968). Ciliogenesis begins with centriole maturation into basal body that nucleates the ciliary axoneme (Avidor-Reiss and Leroux, 2015; Garcia-Gonzalo et al., 2011; Ishikawa and Marshall, 2011; Sorokin, 1962; Williams et al., 2011). Universally conserved 9-fold symmetry of eukaryotic centriole structure templates the 9-fold symmetry of ciliary TZ structure (Lemullois et al., 1988; Sorokin, 1962; 1968). TZ formation is considered to be a one-time structurally deterministic event whose completion precedes, and is required for, formation of the rest of the ciliary axoneme (Avidor-Reiss et al., 2017; Ishikawa and Marshall, 2011; Nechipurenko et al., 2017).

Although a ring of 9 dMT is the most common axonemal pattern of cilia and flagella, TZs of certain protozoans, insect sperm, and nematodes show structural variations containing 3, 6, 8, 10 and 12 dMTs (Dallai et al., 1996; Prensier et al., 1980; Reger and Florendo, 1970; Ross, 1967; Schrevel and Besse, 1975; van Deurs, 1972). Variations in TZ ultrastructure, number of inner sMTs and dMTs, and assembly exist even among cilia with a more conventional 9 dMT axonemal structure (Czarnecki and Shah, 2012; Wiegering et al., 2018). Are cilia with TZ structure comprised of other than nine dMTs formed by the same ciliogenic program as those with 9+0 dMT? Once formed, is the TZ committed to its structure or can the TZ be remodeled to accommodate changing internal and/or external conditions? Surprisingly, no evidence exists on whether TZ with nine-fold radial symmetry of 9 dMTs can change to a TZ structure with less than 9 dMTs, or *vice versa*. Despite the importance of TZ structure to human health (Anand and Khanna, 2012; Bruel et al., 2017; Jauregui et al., 2008; Lambacher et al., 2016; Lu et al., 2017; Williams et al., 2011), our understanding of TZ structural specialization remains incomplete.

We applied transmission electron microscopy (TEM) and electron tomography (ET) in a cross-sectional study to characterize TZ structure in select sensory cilia of a model metazoan *Caenorhabditis elegans*. We chose two types of sensory cilia: <u>ce</u>phalic <u>m</u>ale (CEM) and <u>i</u>nner <u>l</u>abial-<u>2</u> <u>q</u>uadrant (IL2Q), located at dendritic tips of the corresponding sensory neurons located in the head of *C. elegans* (Supplemental Figure 1B). We chose CEM neurons because they have an extended maturation period that coincides with male sexual maturation (Sulston and Horvitz, 1977; White et al., 1986). We hypothesized that such a long developmental window would allow us to capture substeps of ciliogenesis that would otherwise be missed in faster-developing cilia. We chose IL2Q cilia because their TZ have less than 9 dMT TZ structure in wild-type animals (Doroquez et al., 2014; Perkins et al., 1986; Ward et al., 1975). We therefore hypothesized that either IL2Q ciliogenesis includes developmental steps that are incongruent with the current model of ciliogenesis, or that IL2Q cilia undergo post-assembly structural remodeling.

We describe several novel phenomena of TZ structure. First, the number of TZ dMTs in CEM and IL2Q cilia is not static during larval development of *C. elegans*. In developing CEM cilia, the TZ is formed asynchronously building up from 6 to 9 dMTs, while its ciliary axoneme is extended prior to TZ structural completion. IL2Q TZ remodels from an outer 9 to 6 dMT structure. We show that both CEM and IL2Q cilia have structurally dynamic TZs. Our results thus extend the current model of ciliogenesis to reflect structural plasticity of metazoan TZ.

## Results

### Non-canonical 6 dMT transition zone structure of IL2Q cilia is derived by post-assembly remodeling of the canonical 9 dMT structure

Sensory cilia of adult IL2 neurons have a structurally non-canonical transition zone (TZ) that consists of less than nine doublet microtubules (dMTs) (Doroquez et al., 2014; Perkins et al., 1986; Ward et al., 1975). To understand how this non-canonical structure develops, we applied serial section transmission electron microscopy (ssTEM) and electron tomography (ET) approaches to characterize IL2 cilia ultrastructure at three larval stages (L2, L3, L4) preceding adulthood (Supplemental Figure 1A).

Overall, we observed a decreasing trend in dMT number of IL2Q TZ during the developmental period from L2 to adulthood (Figure 1A, B; Supplemental Table 1). In L2 larval males, the IL2Q TZ is comprised of a central cylinder surrounded by 8.0±1.2 dMTs. Because we observed 9 dMT TZs in IL2Q cilia of *both* L2 males and L2 hermaphrodites, we conclude that it is not a sex-specific TZ architecture. In L3 and L4 -stage larval males, the IL2Q TZ contains 7.0±2.0 and 7.5±0.7 dMTs, respectively. In young adults, IL2Q TZ has 6.0±1.2 dMTs on average (Figure 1B). From this we infer that the IL2Q TZ is initially formed before the L2 stage with a canonical structure of 9 dMT. During subsequent larval development, IL2Q ciliary ultrastructure is remodeled to a non-canonical 6 dMT TZ structure seen in adult males (Figure 1) and in adult hermaphrodites (Doroquez et al., 2014; Perkins et al., 1986; Ward et al., 1975). Given the variability of TZ dMT number at each larval stage (Figure 1B), we infer that the timing and extent of this remodeling process is asynchronous.

**Figure 1.**
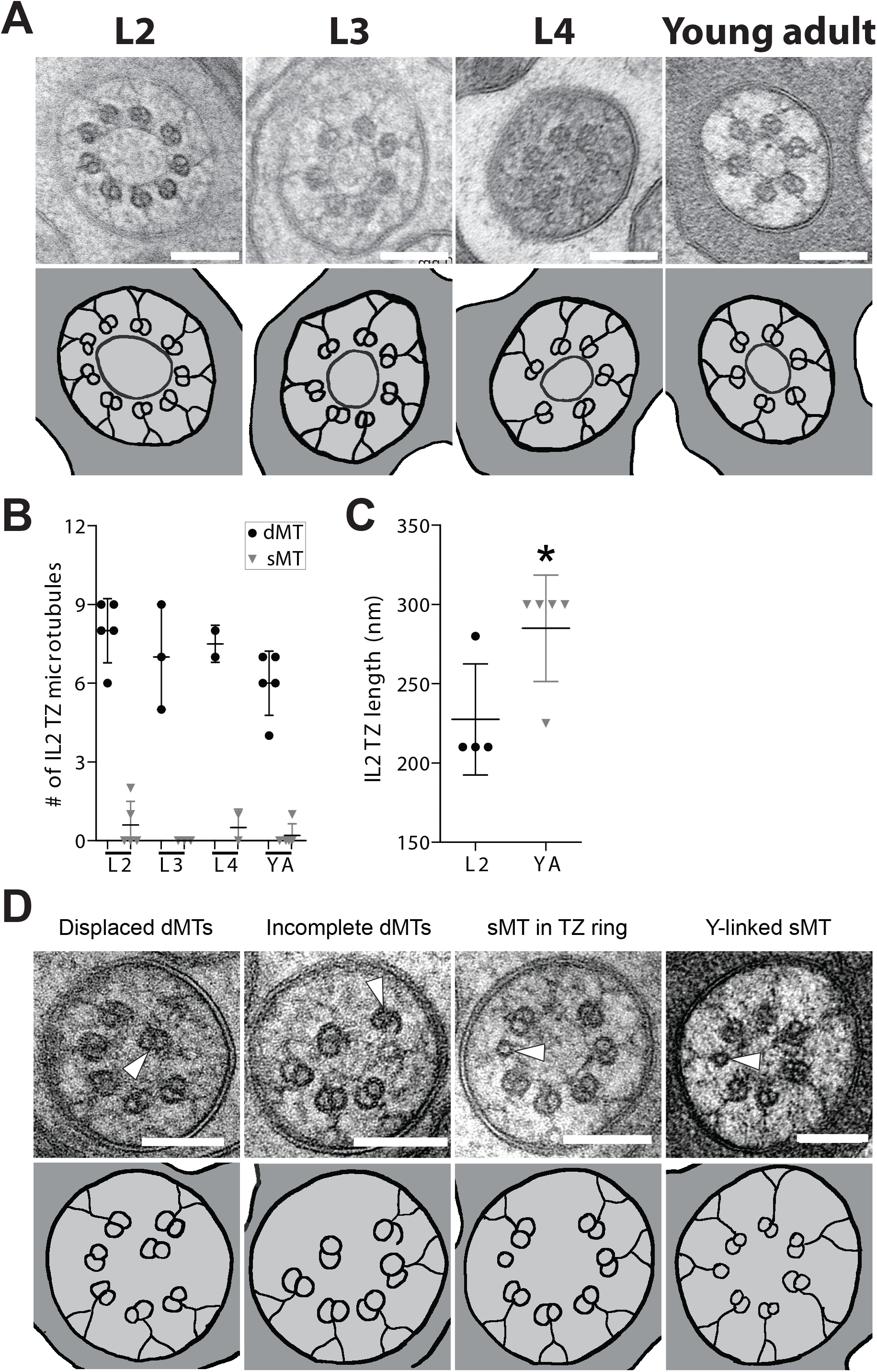
TZ remodeling in IL2Q cilia. **(A)** TEM images (upper row) and corresponding tracings (lower row) of IL2Q cilia TZ in cross-section at L2, L3, L4 and young adult (YA) age animals. L2 image is from a hermaphrodite; L3, L4 and YA from males. Scale bar is 100nm. **(B)** Quantification of the outer dMT and sMTs in IL2Q cilia TZ of L2, L3, L4 and YA males. Each circle (dMTs) and triangle (sMTs) represents measurements from an individual TZ in a different neuron. Errors are SD. The differences between age groups were not significant for either dMT or sMT counts when using the Kruskal-Wallis test with Dunn’s post-hoc correction. See Supplemental Table 1. **(C)** Comparison of IL2Q TZ lengths in L2 larval and young adult (YA) males. Each symbol represents measurements from an individual TZ in a different neuron. Errors are SD. * indicates that marked datasets are different at p<0.05 based on pairwise comparison (Mann-Whitney test). **(D)** TEM images (upper row) and corresponding tracings (lower row) of select cross-sections of larval IL2Q TZs. Arrowheads indicate displaced dMT, A-tubule with hook-like appendage (incomplete dMT), outer sMT without Y link or outer sMT with Y link. Scale bar is 100nm.

At all developmental time points examined, we observed deviations from the stereotypic TZ ultrastructure of circular array of 9 dMTs with Y-shaped membrane tethers (Figure 1D). Occasionally we observed dMTs that are missing Y-links and are displaced towards the center of TZ cylinder. We also observed singlet microtubules (sMTs) with and without Y-links and with hook-like appendages around the central cylinder (herein referred to as “outer sMTs”), where only Y-linked dMTs should be. We infer that the outer sMTs with hook-like appendages are A-tubules with unraveled B-tubules, intermediate forms of microtubules restructuring from AB dMTs to A singlets. Similarly, we infer that outer Y-link-containing outer sMTs are A-tubule remnants of former outer dMTs that completed disassembly of the B-tubule. Because these A-tubule sMTs with hook-like B-tubule remnants are predominately connected at the outer A-B junction (Figure 1D, Figure 3, insert), we infer that of B-tubule disassembly starts at the inner A-B junction and ends at the outer A-B junction. Disassembly of select TZ dMTs notwithstanding, overall length of IL2Q TZ increased from 228±35 nm to 285±34 nm during developmental period from L2 to adulthood (Figure 1C).

Altogether, our TEM and ET findings indicate that the non-canonical 6 dMT TZ structure of IL2Q cilia is derived via remodeling of the canonical 9 dMT TZ structure. This remodeling occurs during larval development and involves asynchronous disassembly of several dMTs. The dMT disassembly process includes unraveling of B-tubules, loss of membrane-tethering Y-links and, eventual disappearance of A-tubules. This structural remodeling reduces the total number of outer microtubules in the TZ structure (Supplemental Figure 1C), but preserves their circular arrangement and the overall length of TZ segment of IL2Q ciliary axoneme (Figure 3).

### CEM ciliogenesis includes asynchronous formation of dMTs via B-tubule synthesis and axonemal extension prior to structural completion of the TZ

In adult males, the CEM cilium TZ is composed of a canonical circle of 9 dMTs that extend as doublets that splay into discrete A-tubule and B-tubule singlets in the middle portion of the axoneme (Silva et al., 2017). Despite being born early in embryogenesis, CEM neurons do not acquire sensory ability until the male reaches adulthood (Chasnov et al., 2007; Silva et al., 2017; Sulston et al., 1983; Wang et al., 2014; White et al., 2007). To test whether the delay in CEM sensory ability is caused by delayed ciliogenesis, we applied ssTEM and ET approaches to characterize CEM cilium ultrastructure at three larval stages (L2, L3, L4) preceding adulthood.

We observed progressive incremental increase in CEM TZ dMT number during the developmental period from L2 to adulthood (Figure 2A, C; Supplemental Table 1). In L2 larval males, the CEM cilium TZ is comprised of circularly arranged mixture of 5.3±1.7 dMTs and 3.0±2.2sMTs, some with hook-like appendages (Figure 2A, Supplemental Figure 1E). Many, but not all, dMTs and sMTs have partial or complete Y-link tethers to the ciliary membrane. The central cylinder is not as pronounced as in IL2Q cilia at this or any of the subsequent ages. In L3 larval males, the CEM TZ has a circular arrangement of 6.8±1.3 dMTs and 0.25±0.5 sMTs. Both dMTs and sMTs have partial or complete membrane tethers. In L4 larval males, the CEM TZ contains 7.4±1.4 dMTs and 0.5±0.8 sMTs, most with distinguishable Y-link attachments to the cilia membrane. In young adults, the CEM TZ contains 8.5±0.7 dMTs arranged in a circle surrounding 4±2 inner singlet microtubules. The inner sMTs are smaller in diameter than outer sMTs seen in larval stages, suggesting a different number of protofilaments in inner and outer sMTs. Outer sMTs are very rare in CEM TZ of young adult males (0.1±0.3). All TZ dMTs have Y-link attachments to the ciliary membrane.

**Figure 2.**
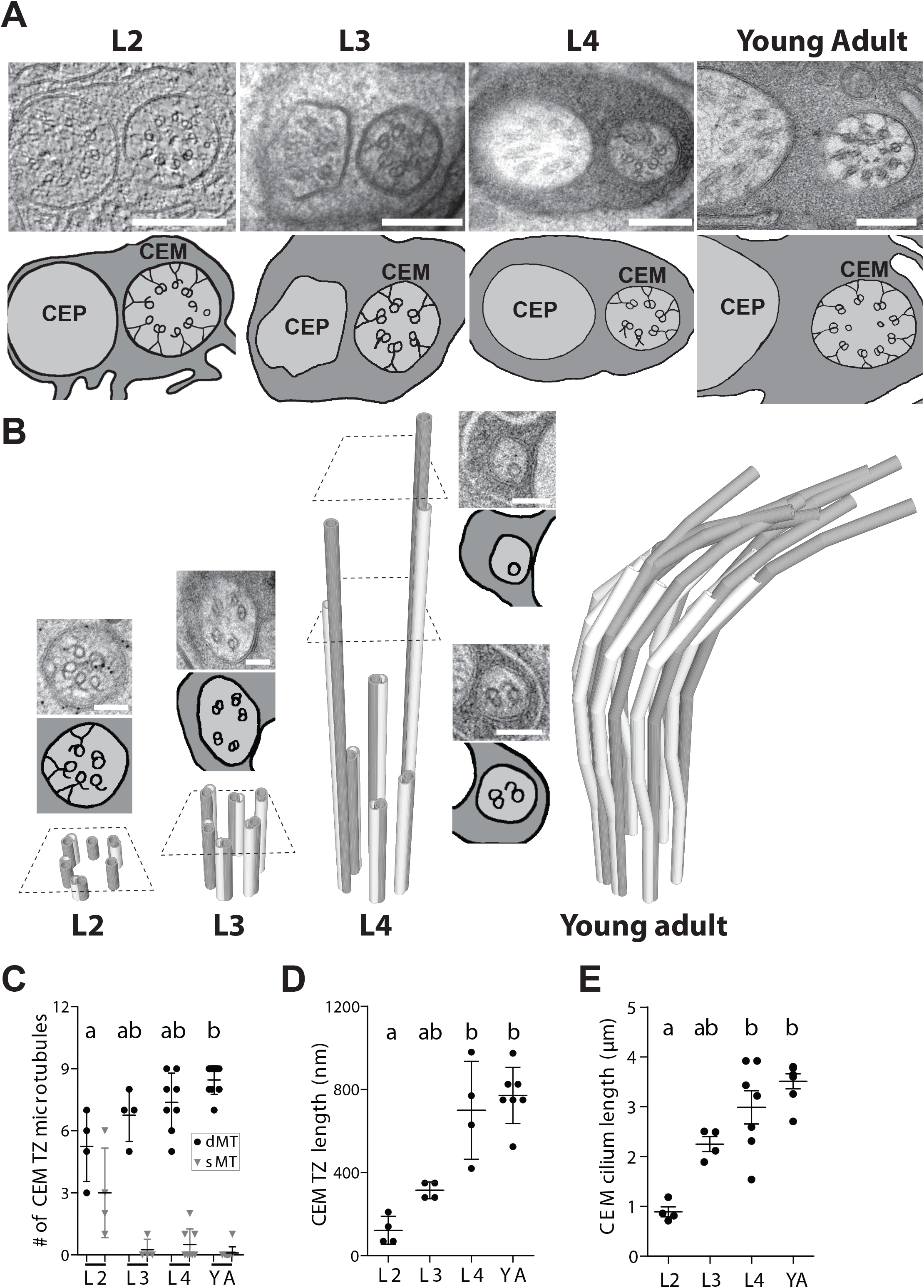
TZ maturation and axonemal extension in CEM ciliogenesis. **(A)** TEM images (upper row) and corresponding tracings (lower row) of CEM cilia TZ in cross-section at L2, L3, L4 larvae and young adult (YA) age males. Scale bar is 100nm. **(B)** Model of growing CEM axoneme anterior of the TZ. Quantity, length and ultrastructure of axonemal MTs is based on serial reconstruction of CEM cilia in L2, L3, L4 and young adult (YA) males. TEM images and corresponding scales show cross-sections of the corresponding CEM axoneme at the focal plane indicated by dashed line. Scale bar is 100nm. **(C, D, E)** Quantification of the outer dMTs and sMTs **(C)**, TZ lengths **(D)** and cilia length **(E)** in CEM cilia of L2, L3, L4 larvae and young adult (YA) males. Each symbol represents measurements from an individual TZ in a different neuron. Errors are SD. Datasets that do not share a common letter are different at *p*<0.005 based on Kruskal-Wallis test with Dunn’s post-hoc correction. See Supplemental Table 1.

Our observations suggest that structural maturation of the CEM TZ includes progressive asynchronous assembly of outer dMTs during L2-L4 larval development to arrive at a canonical 9 dMT TZ structure in the mature adult CEM cilium. The total number of outer microtubules does not change in the CEM TZ during the L2-YA development (Supplemental Figure 1D); but what *does* change, is the inversely proportionate numbers of outer dMTs and sMTs (Figure 2C). This and the presence of outer dMT intermediates (sMTs with hook-like attachments) in larval, but not adult, males, suggest that structural maturation of the CEM TZ occurs via assembly of B-tubules onto to the existing peripheral A-tubule singlets. Because A-tubule sMTs with hook-like appendages are predominately connected at the outer A-B junction (Figure 2A, Figure 3 insert), we infer that of B-tubule assembly begins at the outer A-B junction and ends at the inner A-B junction. Overall length of maturing CEM TZ increases from 122±67nm to 771±135nm during the maturation period from L2 to adulthood (Figure 2D, Supplemental Table 1).

**Figure 3.**
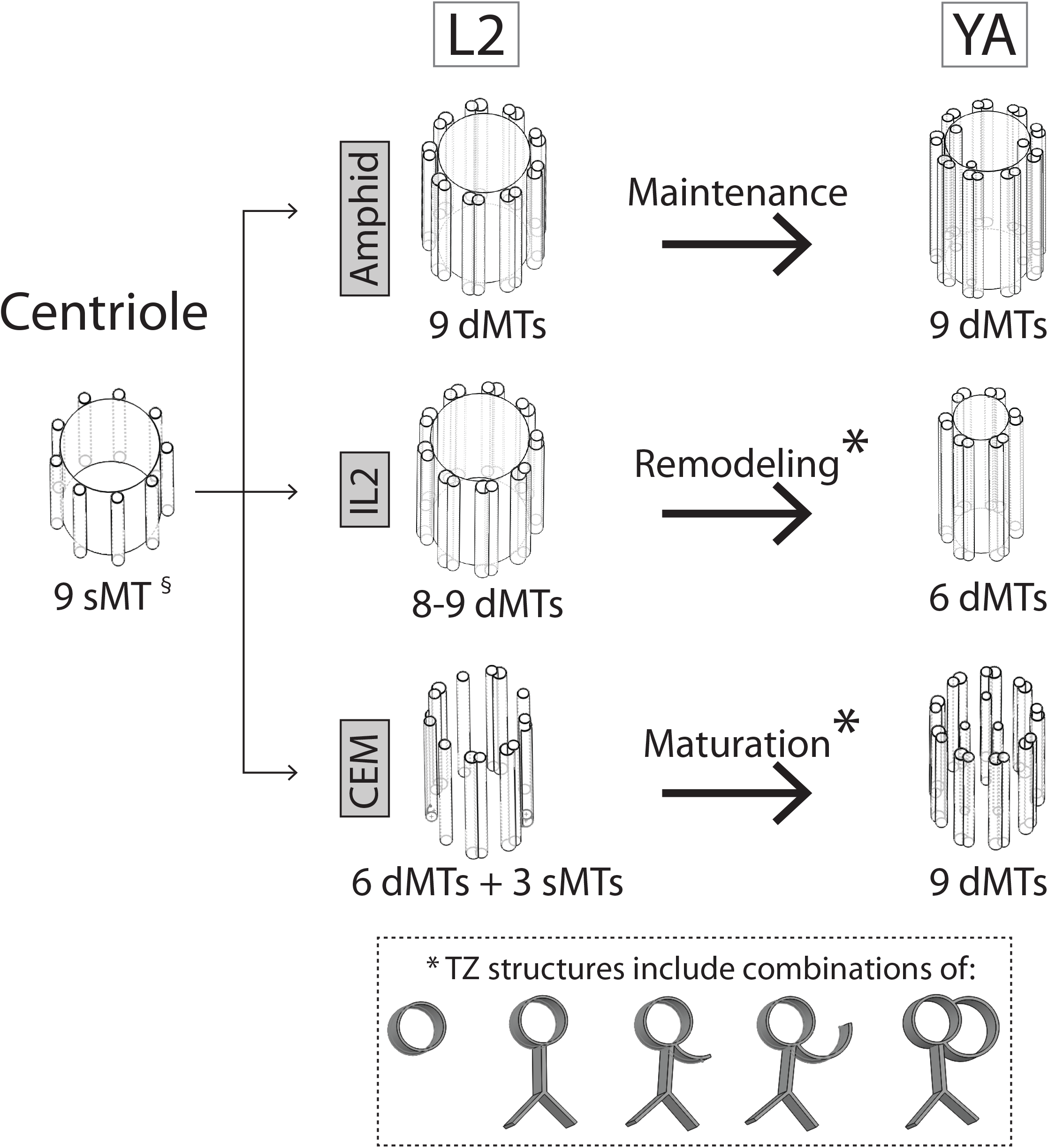
Summary model of age- and cell-specific TZ structural changes. 9-fold symmetry of the centriole templates 9-fold TZ symmetry of sensory cilia in *C. elegans* larvae. Centriolar structure is described in (Pelletier et al., 2006). “Amphid” refers to amphid channel cilia whose early ciliogenesis steps are described by (Nechipurenko et al., 2017; Serwas et al., 2017). Inset shows model of axonemal microtubule structures seen in TZ of IL2 and CEM cilia during L2-YA temporal window. YA = young adult.

Elongation of the CEM ciliary axoneme past the TZ occurs concurrently with TZ maturation (Figure 2B, E; Supplemental Table 1). In L2, L3, and L4 larval males, CEM cilium length is 0.9±0.2μm, 2.3±0.3μm and 3.0±0.9μm, respectively. The lengths of individual axonemal dMT and sMT are increasingly more variable from L2 to L4 (Figure 2B). Given this variability, we conclude that extension of axonemal microtubules in the growing CEM cilium is asynchronous. Examination of the CEM axoneme from distal to proximal revealed that the leading edge of longest axonemal microtubules is an A-tubule singlet, which becomes an A-tubule with hook-like appendage, which becomes a complete dMT at the cilia base (Figure 2A, B). Similar to what was observed in CEM TZ, hook-like appendages are attached to A-tubules at the outer A-B junction (Figure 2B). None of the axonemal dMTs in CEM cilia of L4 males show splayed A-tubule and B-tubule singlet architecture characteristic of adult CEM cilia (Silva et al., 2017).

Despite the incomplete axonemal ultrastructure, CEM cilium length in L4 larval males is comparable to that of adults (Figure 2E), although lacking its stereotypic outward curvature (Figure 2B) (Silva et al., 2017). CEM cilium length in young adults (YA) is 3.5±0.4μm. Measurement via serial section TEM are consistent with our previous measurements based on GFP-tagged axonemal tubulin reporter TBB-4 (Morsci and Barr, 2011; Silva et al., 2017). In contrast to variability of individual axonemal MTs of larval males, in mature adult males all axonemal dMTs run the length of CEM cilium and show specialized splayed ultrastructure with discrete A-tubule and B-tubule singlets in the middle region of the axoneme (Silva et al., 2017).

In summary, the CEM ciliary axoneme is assembled progressively but asynchronously during larval development. As a result, the number of TZ dMTs observed in early larvae does not match the number observed in adult CEM cilia. Our observations suggest that A- and B-tubules of the axonemal dMTs extend asynchronously, with the A-tubule extending first, followed by the assembly of the B-tubule. Surprisingly, CEM cilium length is established by extension of just few microtubules *while TZ* is structurally incomplete. Ultrastructurally, the CEM cilium does not reach its canonical 9 dMTs TZ arrangement and specialized splayed axonemal ultrastructure until sexual maturation and adulthood, more than 72 hours after the onset of its ciliogenesis.

## Discussion

Our observations in CEM and IL2 ciliated sensory neurons of *C. elegans* illustrate that the ciliary TZ can be structurally plastic and undergo ultrastructural architectural changes as part of ciliogenesis (CEM neuron) or post-assembly remodeling (IL2Q neuron) program in wild type animals (Figure 3).

### TZ maturation and remodeling occur via B-tubule assembly and disassembly

Nine-fold symmetry of *C. elegans* sensory cilia structure is templated by a centriole-derived basal body (Nechipurenko et al., 2017; Serwas et al., 2017). Centriolar maturation to basal body includes asynchronous addition of B-tubules to generate AB microtubule doublets in a process that generates structural dMT intermediates (Nechipurenko et al., 2017). Once formed, 9 dMT TZ structure of amphid cilia is maintained throughout larval development into adulthood (Figure 3).

We extend these findings to show that TZ structural maturation during CEM ciliogenesis includes asynchronous assembly of axonemal dMTs primarily via synthesis of B-tubule attachments onto the outer sMTs. The B-tubule of the axonemal dMT was previously shown as a physical substrate of anterograde intraflagellar transport and post-translational modifications in cilia and flagella (Lechtreck and Geimer, 2000; O’Hagan et al., 2011; Redeker et al., 2005; Stepanek and Pigino, 2016). Through its ultrastructure, tubulin composition and post-translational modifications, the B-tubule of the dMT regulates cell-specific coordination of intraflagellar transport motors and, as such, enabling flagellar motility and mediating specialization of ciliary function (Hong et al., 2018; Silva et al., 2017; Suryavanshi et al., 2010). The timing of B-tubule assembly in CEM TZ maturation is asynchronous and temporally stretched out from L2 to young adulthood. We conclude that, the rate of B-tubule synthesis determines the rate of CEM TZ structural maturation and thus the extent of its structural deviation from the canonical 9 dMT architecture. We show that B-tubule synthesis is the mechanism of structural maturation of CEM cilium TZ (Figure 3).

In contrast to what is observed in *C. elegans* amphid ciliogenesis (Nechipurenko et al., 2017; Serwas et al., 2017), we find that CEM TZ structural completion is not a discrete prerequisite step that precedes the rest of axoneme extension. Here we show that, in CEM cilia, outer sMTs and dMTs extend into the axoneme proper *despite* the incomplete TZ structure, with as little as 4 dMT (at L2 stage) to 6 dMT (at L4 stage). This, to our knowledge, is first published evidence that TZ maturation occurs concurrently to axonemal formation and that TZ structural completeness is not required for ciliary axonemal extension. We also show that a growing cilium at near-full length is not necessarily structurally complete nor functionally competent.

Our observations of 9 dMT TZ structure in IL2 cilia of L2 stage larval males suggest that IL2Q TZ formation occurs *before* the L2 stage by the same developmental program and within the same temporal window as in amphid neurons (Nechipurenko et al., 2017; Serwas et al., 2017; Sulston et al., 1983). Unlike ciliated amphid neurons, IL2Q cilia begin to remodel to a non-canonical 6-fold symmetry by selective disassembly of select outer dMTs starting around L2-L3 stage (Figure 1A, B, Figure 3, Supplemental Figure 1A). This remodeling-associated dMT disassembly involves the same structural sMT-dMT intermediates as TZ assembly: A-tubule singlets with hook-like appendages connected at the outer A-B junction and lone Y-linked A-tubule singlets (Figure 1D, Figure 3 inset). The reduction in total number of outer MTs indicates that A-tubule disassembly eventually follows the B-tubule disassembly. Because the IL2Q cilium in young adult male can have as many as 7 or as few as 4 dMTs, we speculate that the *temporal endpoint* of this remodeling is beyond early adulthood; the *structural endpoint* is unknown. This IL2Q TZ remodeling is genetically controlled, as evidenced by select ciliogenesis mutants that show ectopic 9 dMT TZ structure in IL2 cilia in adult animals (Perkins et al., 1986).

### Wild-type CEM and IL2 TZs have incomplete dMTs and sMTs with Y-links

Defects in cilia formation or function are implicated in a variety of human diseases called ciliopathies (Reiter and Leroux, 2017). Phenotypic characterization of cilia in disease contexts extensively relies on the canonical structure of a ring of outer 9 dMT as reference for normal healthy cilium. As a consequence, the presence incomplete dMTs (A-tubules with hook-like appendages) in ciliary axoneme may be characterized as a degenerative phenotype. Our observations show that presence of incomplete dMTs, and Y-linked sMTs in the TZ may also be normal characteristics of an axonemal maturation step of ciliogenesis or evidence of post-assembly structural remodeling. These findings are consistent with previous observations of sMTs with hook-like appendages in developing amphid cilia TZs in wild-type *C. elegans* (Nechipurenko et al., 2017). This suggests that ciliary mutant phenotypes may be developmental or degenerative. While degenerative ciliary phenotypes may become progressively worse with time, a developmental phenotype *may improve with time* in certain mutant backgrounds. A recent publication may already have showed evidence of this, describing structural and functional time-dependent *recovery* of sensory cilia from young adulthood to middle age in *C. elegans* hypomorphic mutants previously known for axonemal structural defects (Barr, 2016; Cornils et al., 2016). Similarly, *Drosophila* mutants that lack TZ-localizing MKS proteins and show axonemal structural defects in larval development *recover* in adulthood (Pratt et al., 2016).

In both CEM and IL2Q cilia, TZs structural remodeling is genetically controlled, since some mutants have ectopic 9 dMT TZ in adult IL2 cilia (Perkins et al., 1986) or less than 9 dMT ectopic TZ in adult CEM cilia (Wang et al., 2015). Further investigation into the mechanisms of these axonemal remodeling processes raises exciting possibilities of being able to apply them for therapeutic purposes, in cases of ciliopathies caused by structural TZ defects. This has profound implications to human health and all the ciliary diseases that are characterized by structural defects of ciliary axoneme.

### The role of TZ structural plasticity in ciliary function and animal development

We propose that TZ structural changes reported in this work are supporting the emergence (CEM cilium) or modification (IL2Q cilium) of ciliary sensory function.

CEM neurons are male-specific chemosensory neurons that detect secreted pheromones of the opposite sex (Narayan et al., 2016; Srinivasan et al., 2008) and release extracellular vesicles that regulate male mating behavior (Silva et al., 2017; Wang et al., 2014). CEM neurons are born, migrate and extend dendrites in the same temporal window of embryogenesis as other ciliated head neurons, including amphids (Supplemental Figure 1A) (Singhal and Shaham, 2017; Ward et al., 1975). Amphid channel cilia complete their 9 dMT TZ structure within 2 hours of neurogenesis and maintain TZ structure through larval development into adulthood (Nechipurenko et al., 2017; Perkins et al., 1986; Serwas et al., 2017). These neurons are functional by the time L1 larva hatches from its egg. Unlike the amphid neurons, CEM neurons *neither* complete their axonemal ultrastructure *nor* become functionally competent until the male reaches adulthood, more than 72 hours post neurogenesis (Figure 2A, B; Supplemental Figure 1A) (Chasnov et al., 2007; Narayan et al., 2016; Silva et al., 2017; Wang et al., 2014; 2015; White et al., 2007). p38 MAPK *pmk-1* mutants that fail to complete the assembly of 9 dMTs CEM TZ are defective in mating behaviors (Wang et al., 2015). Here we show that the latent onset of CEM sensory competence is, at least in part, contributed to by latent ultrastructural maturation of its ciliary axoneme. We speculate that CEM ciliogenesis is temporally tuned to the development of the whole organism, such that its structural completion is timed to occur no sooner than the age at which CEM sensory ligands, sex pheromones, can be acted upon by the male in age-appropriate manner.

IL2Q TZ structural differences between larval and adult animals may also confer stage-specific functional differences. IL2 neurons have previously been shown to be structurally dynamic and plastic in response to changing developmental cues. During L1 transformation into dauer, an alternative developmental stage triggered by harsh environmental conditions, IL2Q neurons undergo dramatic dendritic arborization and axonal remodeling (Schroeder et al., 2013). Functionally, IL2 cilia are required for the nictation behavior of dauer larvae (Lee et al., 2011). In adult hermaphrodites, IL2 neurons are required and sufficient for stimulatory action of ethanol on locomotion (Johnson et al., 2017). This effect is blocked in several ciliary mutants (Dr. J. Barclay, pers. comm.), indicating that ethanol stimulation is probably mediated by IL2 cilia function. We speculate that IL2Q cilia may have one function in larvae, which requires 9 dMT TZ structure, and another function in adults, which requires 6 dMT TZ structure. Unfortunately, sensory ligand of IL2 cilia is yet to be discovered. Once IL2 sensory stimuli are identified, it would be interesting to compare the IL2 cilia sensory ability in larvae (9 dMT TZ) to adults (6 dMT TZ) and mutant adults with ectopic 9 dMT TZ structure in adulthood (Perkins et al., 1986). These future studies will reveal whether remodeling of 9 dMT into 6 dMT structure contributes functional reprogramming of IL2 cilia for a different sensory modality.

The CEM and IL2 neurons are the only ciliated neurons in the brain of the worm that shed and release ciliary extracellular vesicles (EVs) (Wang et al., 2014; Wang et al., 2015; Wang and Barr 2016). EVs are nano-communication devices that cells shed and release to influence the behavior of other cells, tissues, or even organisms (Maas et al., 2017). EVs may be beneficial or toxic, depending on their cargo. Very little is known about EV cargo sorting, formation, or function, largely because their tiny size (100nm) escapes detection by light microscopy. We previously showed that ciliary EV shedding and release is regulated by the intraflagellar transport machinery, microtubule glutamylation, and a cell-specific ciliary kinesin KLP-6 and α-tubulin TBA-6 (Wang et al., 2014; Wang et al., 2015; Silva et al., 2017; O’Hagan et al., 2017). We propose that the cell-type specific structural plasticity of the CEM and IL2Q ciliary TZ gate may influence ciliary EV biogenesis including cargo sorting, shedding, and/or environmental release (Wang and Barr 2018). Consistent with this hypothesis is the finding that TZs isolated from *Chlamydomonas* flagella are enriched for endosomal sorting complex required for transport (ESCRT) proteins involved in ciliary EV release (Diener et al., 2015; Long et al., 2016).

The ciliary TZ acts as a selective gate that regulates protein entrance and exit between the cell and cilium (Garcia-Gonzalo and Reiter, 2017). The discovery of TZ structural plasticity raises some intriguing questions: What mechanisms drive TZ remodeling and maturation? How does molecular composition of the cilium change during TZ remodeling and maturation? Does changing TZ structure affect properties and composition of the ciliary membrane and of ciliary-derived extracellular vesicles? Is the selectivity of the ciliary gate impacted by the structural conversion of the TZ from the canonical 9 dMT structure to a non-canonical one? Do fewer Y-linked MTs make for a more permissible ciliary gate? Our studies in *C. elegans* reveal novel additions for the established view of ciliogenesis and determination of TZ structure, and will continue to provide new ways to investigate cilium biology.

## Conclusions

Our studies reveal that metazoan TZ is remarkably plastic even in the context of terminal post-mitotic differentiated cells like neurons. We show that a non-canonical 6 dMT TZ structure can be derived from a canonical 9 dMT TZ and *vice versa*; both of the processes mediated by B-tubule structural dynamics. We propose that TZ structural remodeling may mediate the emergence or change in ciliary function. The current model of ciliogenesis needs to be expanded to include the structurally diverse, plastic and dynamic process of TZ maturation, maintenance and remodeling.

## Materials and Methods

### *C. elegans* strains

CB1490 *him-5(e1490)* V

PT443 *myIs1* [PKD-2 ∷GFP + Punc-122∷gfp] *pkd-2(sy606)* IV; *him-5(e1490)* V

Supplemental Figure 1B:

PT2102 *pha-1(e2123)* III; *him-5(e1490)* V; myEx686 [Pklp-6∷GFP∷gKLP-6_3’UTR + pBX]

### High pressure freeze fixation (HPF) and freeze substitution (FS)

Age matched animals (PT443 for L2 stage animals and CB1490 for all other stages) were subjected to high-pressure freeze fixation using HPM10 high-pressure freezing machine (Bal-Tec, Switzerland). Animals were freeze substituted in 2% osmium tetroxide, 0.2% uranyl acetate and 2% water using RMC freeze substitution device (Boeckeler Instruments, Tucson, AZ, USA) (Weimer, 2006). Samples were infiltrated with Embed 812 resin over three days prior to embedding in blocks. L2 males were picked based on coelomocyte positions, which are different in males and hermaphrodites (Sulston and Horvitz, 1977).

### Transmission electron microscopy (TEM)

For TEM, 75nm plastic serial sections were collected on copper slot grids. Sections were post-stained with 2% uranyl acetate in 70% methanol, followed by washing and incubating with aqueous lead citrate. TEM images were captured on either Philips CM10 Transmission Electron Microscope operating at 80kv or JEOL JEM-1400 Transmission Electron Microscope operating at 120kv.

### Electron tomography (ET)

We used a combination of single and dual axis tomograms. Tilt series acquired from either thin (75nm) or thick sections (250nm) using an FEI Technai20 TEM or a JEOL JEM-1400 using SerialEM software with 140 images per tilt axis. These data were Fourier processed using IMOD software (Hall and Rice, 2015; Mastronarde and Held, 2017); tomograms were generated using either the marker-less back projection method (Kremer et al., 1996) or simultaneous iterative reconstruction technique (SIRT) (Wolf et al., 2014).

### Figures and image handling software

Tomograms were viewed using IMOD software (Hall and Rice, 2015; Mastronarde and Held, 2017). FIJI software was used to visualize both tomograms and TEM images. Some of the EM pictures were ‘slice-views’ exported from tomograms. Serial TEM images were stacked, aligned, annotated using the TrakEM2 suite in FIJI (Schindelin et al., 2012). Final figures with TEM images were made using a combination of FIJI, Adobe Photoshop, and Adobe illustrator. Graphics in Figure 2B and Figure 3 were made using SOLIDWORKS™ 3D CAD software.

### Data collection and statistical analysis

TZ length was estimated from the number of cross-section ssTEM images with clearly visible Y-links. Microtubule numbers in CEM and IL2Q TZ were counted based on cross-section TEM images at the posterior-most ends of the TZs, where all the dMTs have Y-links to the ciliary membrane. For differential quantification of microtubule singlets (sMTs) and doublets (dMTs) in Figures 1B and 2C, microtubules with hook-like appendages were considered as singlet microtubules (sMTs). TZ lengths were estimated using serial TEM images. Statistical analysis and graphing was done using GraphPad Prism V5. Pairwise comparison was done using Mann-Whitney test. *, ** and *** indicate significance based on *p* <0.05, <0.005, and<0.0005, accordingly. When comparing more than two groups, the Kruskal-Wallis test with Dunn’s post-hoc correction was used, in which case we used distinct letters to distinguish groups with statistically significant difference. *p* values of all tests are listed in Supplemental Table 1. Lowercase “n” indicates the number of cilia, while upper case “N” indicates number of animals.

## Abbreviations

TZ: transition zone; dMT: microtubule doublet; sMT: microtubule singlet; YA: young adult; CEM: cephalic male (neuron); IL2Q: inner labial 2 quadrant (neuron); L2, L3, L4: larval stage 2, 3, 4

---

**Supplemental Figure1.**
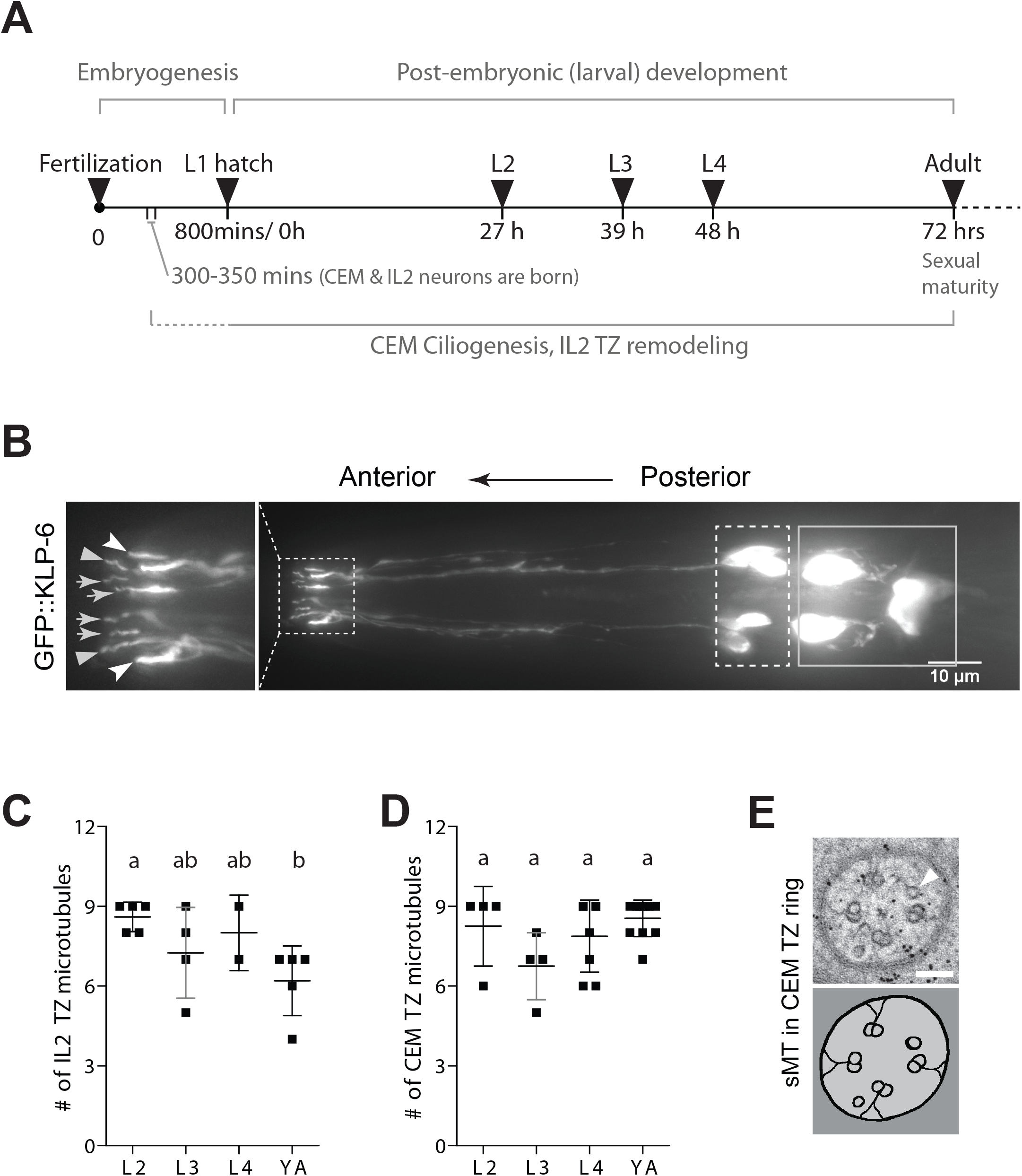
**(A)** Developmental timeline of *C. elegans* at 20°C. *C.elegans* undergoes four larval stages of development prior to reaching adulthood. L1, 2, 3, and 4 indicate successive larval stages of development. Embryogenesis span is counted in minutes (from 0 to 800), larval development in hours, starting from L1 larvae hatching from the egg to the last molt from L4 to adulthood, from 0 to 72 hours, accordingly. The approximate time period during which CEM and IL2 neurons are born, undergo ciliogenesis and remodeling are marked with solid lines. Dashed line indicates uncertainty of the starting points for the respective processes. **(B)** Fluorescent micrograph of the head of a young adult male *C. elegans* expressing a GFP-tagged kinesin-3 KLP-6 in the IL2 and CEM neurons. The IL2 neuron cell bodies are enclosed in dashed line box, and the CEM neuron cell bodies are enclosed in solid line box. Inset shows the IL2 and CEM cilia, located at the anterior ends of the sensory dendrites. Grey arrows point to the IL2Q cilia; grey *arrowheads* point to the IL2L cilia (not studied in this paper). White notched arrowheads point to CEM cilia. Scale bar is 10μm. **(C, D)** Quantification of the total number of outer microtubules in IL2Q (C) and CEM (D) cilia, where sMTs and dMTs are counted equally. Each square represents a measurement from an individual TZ in a different neuron. Errors are SD. Letters above indicate results of statistical comparison, where groups that share a common letter are not significantly different from each other. Groups were compared using Kruskal-Wallis test with Dunn’s post-hoc correction. Significance was called at p <0.05. See Supplemental Table 1 for additional details. **(E)** TEM image of a CEM cilium TZ in cross-section. Arrowhead is pointing at the outer sMT. Scale bar is 100 μm.

**Supplemental Table 1.**
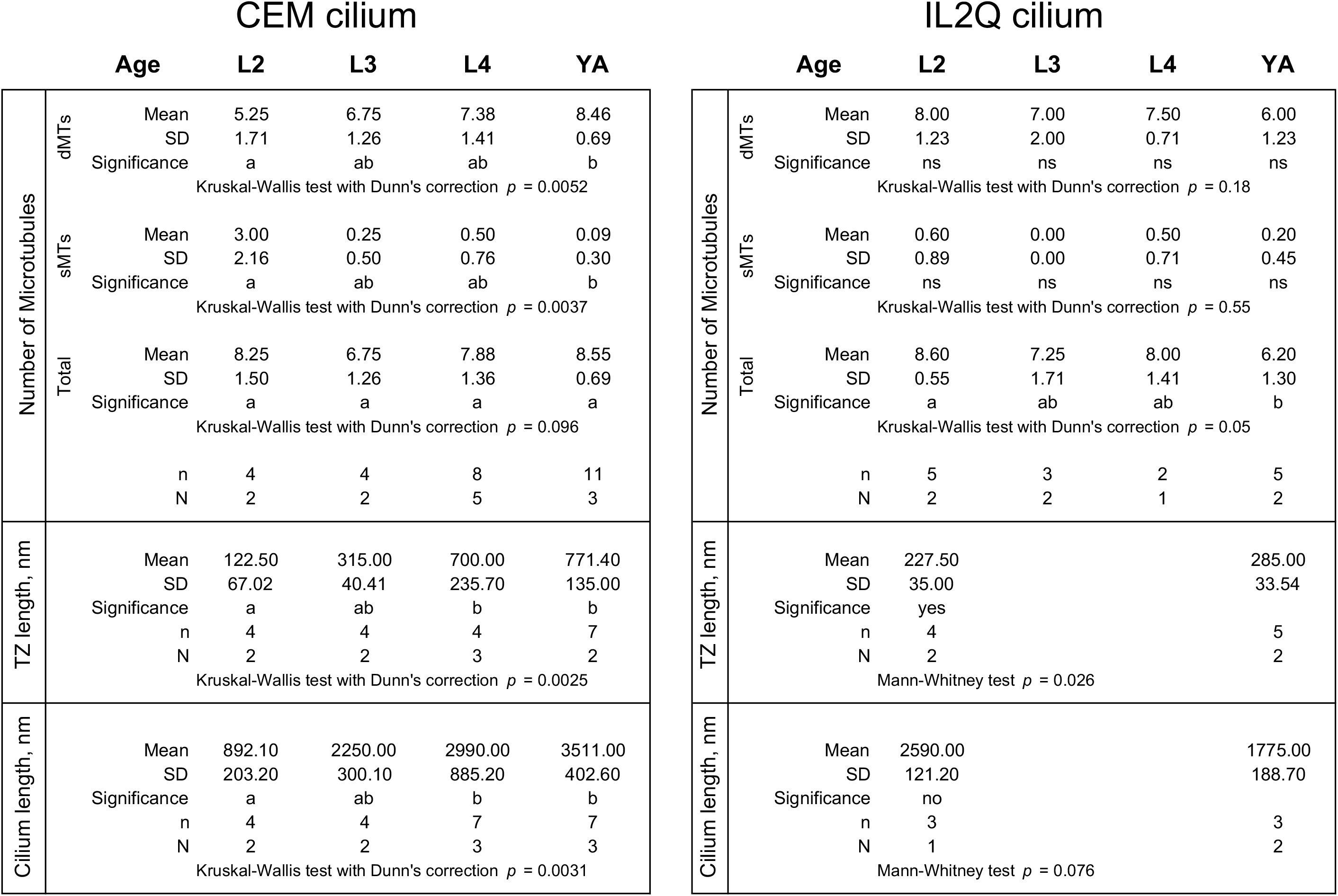
Data from Figures 1, 2, S1 and Statistical analysis. Statistical analysis and graphing was done using GraphPad Prism V5. Pairwise comparison was done using Mann-Whitney test. *, ** and **1 indicate significance based on *p* <0.05, <0.005, and<0.0005, accordingly. When comparing more than two groups, the Kruskal-Wallis test with Dunn’s post-hoc correction was used, in which case we used distinct letters to distinguish groups with statistically significant difference. Lowercase “n” indicates the number of cilia, while upper case “N” indicates number of animals.

## Authors’ contributions

Conceptualization: J. S. A., M.S., and M. M. B.; Methodology, investigation, and analysis: J. S. A., M. S., N.S.M., K. Q. N., W. J. R, D. H. H., and M. M. B.; Manuscript draft writing: J. S. A., M. S.; Writing and editing of the submitted manuscript: N. S. M., M. M. B.

## Acknowledgements

We thank Leslie Gunther and Frank Macaluso for help in HPF-FS performed at Einstein, Ed Eng at the New York Structural Biology Center (NYSBC) for help in electron tomography; Gloria Androwski for ongoing outstanding laboratory support; WormBase and WormAtlas for online resources; the Barr lab for discussion and constructive criticisms; Joel Rosenbaum for feedback on the manuscript.

